# Constrained template matching using rejection sampling

**DOI:** 10.1101/2025.06.10.658890

**Authors:** Valentin J. Maurer, Lukas Grunwald, Dante M. Kennes, Lea Dietrich, Jan Kosinski

## Abstract

Identifying macromolecular complexes in situ using cryo-electron tomography remains challenging, with low signal-to-noise ratios, the missing wedge, and crowded backgrounds among the key limiting factors. By integrating prior knowledge on macromolecular localization, such as the preferred orientations of membrane-associated proteins, detection can be improved by constraining searches to biologically feasible orientations. Here we describe rejection sampling, an approach for integrating such constraints at voxel resolution that remains both accurate and computationally efficient. Using synthetic and experimental data, we show that these constraints improve detection, orientational assignment, and discrimination between macromolecules. The resulting picks match the performance of deep-learning methods informed by membrane structure without requiring annotated data. We further apply rejection sampling to ATP synthase on mitochondrial cristae, illustrating how it extends macromolecular detection to the large and highly curved membrane systems that pervade cells.

## I. INTRODUCTION

Cryo-electron tomography (cryo-ET) has become a cornerstone of structural biology, enabling visualization of macromolecular complexes within their native environments at molecular resolution^1–3^. By imaging frozen-hydrated samples at different tilt angles and computationally reconstructing three-dimensional tomograms, cryo-ET provides detailed views into molecular architecture and protein organization in situ^4^. However, automated detection of macromolecular complexes is complicated by low signal-to-noise ratios (SNR), the missing wedge, and contrast transfer function (CTF) modulation inherent to data acquisition, as well as by molecular crowding^5,6^.

Template matching enables macromolecular detection by systematically comparing a reference template representing a specific macromolecule of interest against a tomogram based on cross-correlation^7^. This approach has proven effective for detecting isolated particles in purified samples but has low precision and recall in crowded cellular environments^5,7,8^. However, when prior knowledge of the orientation of macromolecules relative to cellular geometries is available, such as the preferred orientations of membrane-associated proteins, the translational and rotational searches can be constrained to biologically relevant configurations to improve detection^9^.

Such prior knowledge can be modeled through seed points, initial guesses of the position and orientation of macromolecules that can be derived from surface parameterizations of membranes. The standard approach extracts a subtomogram at each seed point to align the template locally, limiting translations to a subregion and orientations to within a predefined angular cone of the seed point normal^10,11^. Extensions of this idea reframe the alignment as a template-matching task on subtomograms over a constrained angular range^12^. Faithfully representing the surface geometry, and thus restricting matches to biologically plausible configurations, requires more seed points as system size and curvature increase. For large and strongly curved systems such as viral envelopes and mitochondrial cristae, the required sampling density becomes computationally prohibitive on commonly available hardware. Subtomogram-based approaches therefore face a tradeoff between the fidelity of the geometric constraints and the computational cost of enforcing them.

Recent deep learning-based approaches avoid this tradeoff by treating the membrane as a continuous surface. MPicker^13^ resolves the curvature explicitly, flattening the membrane and reducing detection to two-dimensional picking on orthogonal slices with EPicker^14^. In contrast, MemBrain-pick^15,16^ forgoes explicit flattening, sampling densities along the surface normal, and passing them to a network that retains three-dimensional context. Such detectors localize membrane proteins effectively, but require annotated positions of the target complex for training. They also have no direct means of determining the full particle orientation without additional processing. For template matching, flattening would likewise simplify the rotational search, but imaging corrections required for accuracy, such as the missing wedge and CTF^7,9^, are tied to the original imaging geometry and do not transfer straightforwardly to the locally deformed space flattening produces. A method that overcomes these limitations must therefore operate in the original tomogram space, where these corrections remain valid, represent curved surfaces at voxel resolution, and determine particle position, identity, and orientation from a reference alone.

To achieve this, we introduce constrained template matching using rejection sampling. Rather than restricting the search to individual subtomograms, this approach enforces the constraints across the original tomogram, enabling surface representation at voxel resolution. Using synthetic and experimental cryo-ET data, we show that these constraints are critical for accurately detecting, orienting, and distinguishing different macromolecules, that they add negligible computational overhead, and that the resulting picks are competitive with deep-learning methods informed by membrane structure. We then outline the application of rejection sampling to ATP synthase in mitochondrial cristae of primary rat neurons.

## II. RESULTS

Rejection sampling casts constrained template matching as a proposal-and-acceptance procedure, in analogy to its Monte Carlo counterpart. Schematically, it follows the unconstrained template-matching procedure over the entire tomogram, computing cross-correlation scores for all template translations and a set of rotations; this approximately uniform grid serves as the proposal. Constraints are encoded by seed points, which can represent any prior knowledge of macromolecular arrangement but, in this work, derive from membrane surfaces. After each rotation, scores that violate the imposed constraints are rejected and can be thought of as biologically implausible. A score is accepted if the translation it describes lies within the acceptance region of a seed point, and if its rotation, in the local coordinate basis of that seed point, deviates from the normal by no more than a defined threshold (Fig. 1).

**Fig. 1.**
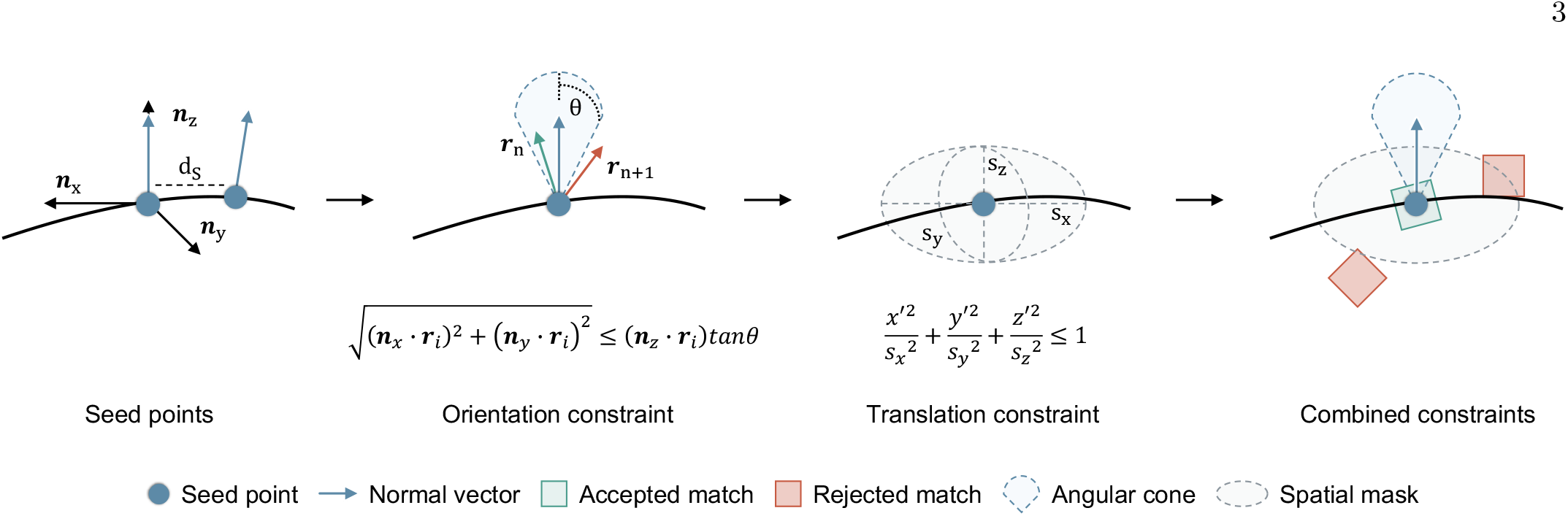
Conceptual depiction of constrained template matching using rejection sampling. A seed point with normal vector ***n*** *≡* ***n***_*z*_ defines the local coordinate system with basis ***n***_*x*_, ***n***_*y*_, ***n***_*z*_. d*s* denotes the average spacing between seed points. The orientational constraint limits template orientations to those where the reference vector ***r***_*i*_ falls within a cone of aperture *θ* around the normal, e.g., ***r***_*n*_. The translational constraint is implemented as an ellipsoid with radii *s*_*x*_, *s*_*y*_, *s*_*z*_ centered at each seed point *µ*_*x*_, *µ*_*y*_, *µ*_*z*_ and aligned along ***n***_*z*_. Only matches satisfying both orientational and translational constraints are retained.

Each seed point thus enforces constraints locally, and the seed points collectively approximate the global constraint imposed by the membrane surface. This approximation improves as the seed point spacing tightens and each acceptance region narrows to just span the gap to the next seed point.

### A. Geometric constraints improve detection and discrimination of macromolecules

To evaluate rejection sampling and the contribution of its constraints, we constructed a synthetic Influenza A virus (IAV) comprising a spherical membrane decorated with hemagglutinin (HA) and neuraminidase (NA) at reported ratios and spacings^17,18^. From this model, we simulated a tomogram^19^ incorporating solvent scattering, missing wedge, CTF modulation, and acquisition noise (Fig. 2a). These data served as ground truth for the comparisons below.

**Fig. 2.**
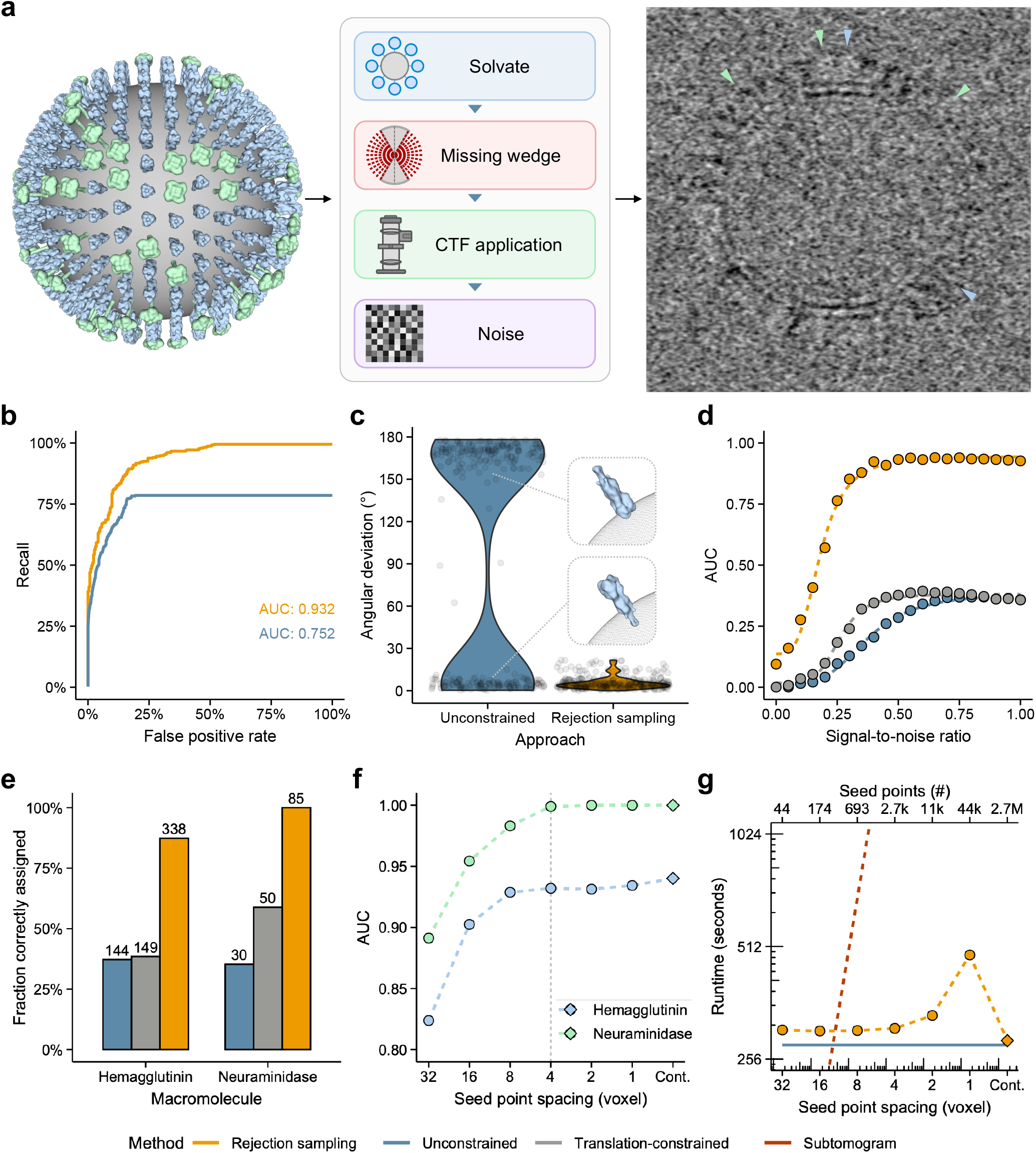
Evaluation of rejection sampling on synthetic cryo-ET data. **a**, Simulating a tomogram from an IAV envelope model consisting of HA (blue) and NA (green), accounting for solvent, missing wedge, CTF, and acquisition noise. Colored triangles indicate HA and NA instances in the simulated tomogram (right). **b**, Receiver operating characteristic (ROC) curves for HA detection at an SNR of 0.65 (top 650 picks used for all experiments). **c**, Angular deviation between matched and ground truth HA orientations at an SNR of 0.65 (*n* = 385 and 304 for rejection sampling and unconstrained, respectively). Violins indicate kernel density estimates, and points indicate individual matches. Given the prevalence of inverted orientations, a match in (**d**)-(**g**) is correct if it lies within 30° of the ground truth orientation, unlike in (**b**), which considers position alone. **d**, AUC for HA detection across SNRs, comparing full constraints (rejection sampling) to translation constraints (translation-constrained) and no constraints (unconstrained). SNR refers to the corresponding value set during data simulation. **e**, Discrimination between macromolecules. A detection was scored as correct if the corresponding template produced a higher cross-correlation score at that position (e.g., the HA template scored higher for HA particles). Numbers are absolute particle counts. **f**, AUC versus seed point spacing for HA and NA. Cont. denotes the continuous-constraint formulation. Dashed line: spacing used in (**b**) and (**c**). **g**, Runtime across methods. Points are measurements, and dotted lines are projections. Unconstrained matching is shown as the baseline. The top axis indicates the seed point count at the given spacing.

As a baseline, unconstrained template matching achieved an area under the curve (AUC) of 0.752 for HA detection (Fig. 2b). Many HA instances were localized correctly, but their orientations were inverted, such that the ectodomain pointed toward the membrane interior in 56% of matches (angular deviation *>* 90°, Fig. 2c). NA detection was more accurate (Fig. S1a) but showed the same orientation failure, in this case for 70% of matches (Fig. S1b). Rejection sampling raised HA detection to an AUC of 0.932 and confined all matches to within the specified 20° cone, with median deviations of 4.4° (HA) and 4.1° (NA) well inside that bound. Thus, correct localization does not guarantee correct orientation without explicit orientational constraints.

Erroneous detections can arise from a nonspecific background, which can be suppressed by masking cross-correlation scores. This is functionally equivalent to rejection sampling without orientational constraints, and we hereafter refer to this as translation-constrained. Comparing all three approaches across a range of SNRs (Fig. 2d), translation-constrained matching improved on unconstrained matching throughout but plateaued at substantially lower AUC than rejection sampling, which maintained robust performance down to an SNR of approximately 0.375. To test whether this gap extends to distinguishing HA from NA, we matched both templates independently and assigned each detection to the higher-scoring template. The same ordering held, with rejection sampling correctly assigning roughly twice as many instances as translation-constrained matching (Fig. 2e). Orientational constraints are thus essential for reliably determining macromolecular orientation and identity.

These experiments used a fixed seed point spacing, but the appropriate spacing is not straightforward in general. It depends on surface curvature (Section XI A) and, for a sphere, simplifies to d*ω* = d*sR*^−1^. This predicted a spacing of approximately 8 voxels for this system, a value we conservatively halved in the experiments above. Detection performance was indeed largely insensitive to spacing below this prediction (Fig. 2f). However, the runtime was not, as a 1-voxel spacing doubled the runtime relative to unconstrained matching (Fig. 2g). This remains far faster than using subtomograms, which scales linearly with seed point count, yet it becomes limiting for larger/curved systems. This can be avoided by defining each voxel within a given distance of the membrane as a seed point and shrinking each acceptance region to that voxel. With this formulation, each constraint can be evaluated independently, thereby greatly reducing runtime. In fact, it is essentially that of template matching alone (Fig. 2g). We develop this formulation in the next section.

### B. Continuous constraints enable accurate detection in complex membrane systems

Thylakoid membranes house the photosynthetic machinery of plant chloroplasts, organized into tightly stacked grana interconnected by stromal lamellae^20^. Within grana, adjacent membrane sheets are separated by a stromal gap of only 3.2 nm^21^ and harbor a high concentration of Photosystem II (PSII), which catalyzes the photolysis of water^22^. Embedded in this closely stacked, high-contrast membrane system, PSII is a challenging target for template matching, and for the discrete seed point formulation in particular.

On tightly stacked membrane sheets, the translational acceptance bands of seed points from adjacent membranes overlap, assigning multiple normal vectors to the same voxel. This ambiguity undermines a central guarantee of rejection sampling, namely that the orientational constraint provides a strict upper bound on the angular error of accepted matches. We therefore developed a continuous formulation in which the constraints derive from the normal vector field of a signed distance function for the membrane surface, thereby assigning a single normal to every voxel (Fig. 3a). This continuous formulation restores the upper bound and reduces the translational constraint to a single parameter, *h*, the maximum distance to the surface.

**Fig. 3.**
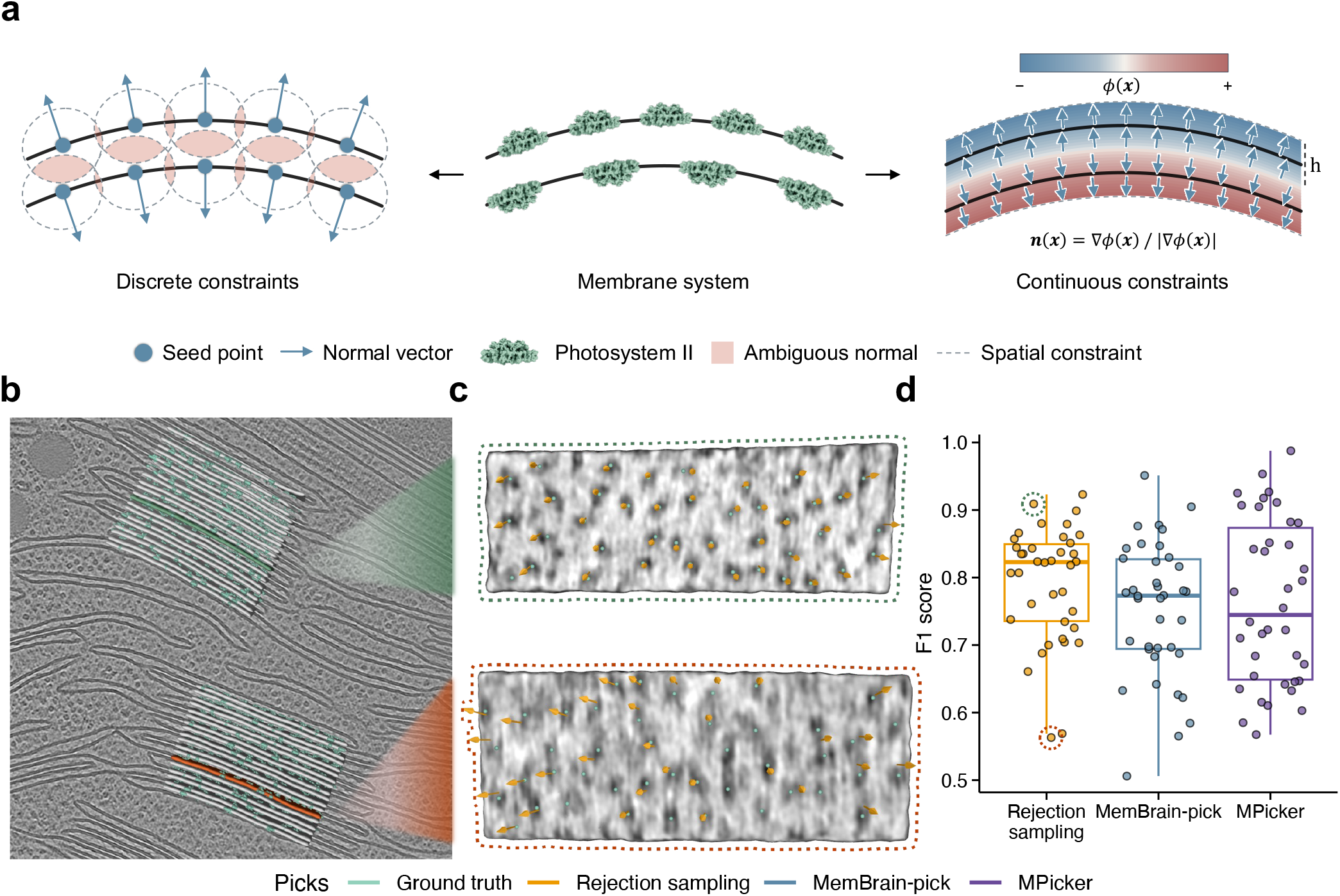
PSII localization on spinach thylakoid membranes. **a**, Discrete seed point constraints produce normal vector ambiguity in tightly stacked membranes due to overlap of their acceptance regions. This is resolved using continuous constraints derived from the signed distance function of the membrane surface. *h* indicates the accepted maximum distance from the surface. **b**, Rendering of spinach chloroplast thylakoid membrane stacks, a subset of which has been segmented and meshed (grey). PSII ground-truth annotations are shown as densities (green). **c**, Representative membrane sheets on which rejection sampling achieved high (top) and low (bottom) F1 scores, with picks, orientations, and ground truth positions indicated. Tomographic densities were projected onto the surface. **d**, F1 score across membrane sheets shown in (**b**, *n* = 38) stratified by method. Colored circles mark membranes shown in (**c**). Boxes correspond to the median and quartiles. Whiskers indicate the furthest point within 1.5*×* interquartile range of the hinges.

Using these continuous constraints, we applied rejection sampling to cryo-ET data of spinach chloroplasts with available PSII ground truth annotations (Fig. 3b)^20^. On membranes with well-resolved PSII densities, picks closely matched the ground truth (Fig. 3c). On lower-contrast membranes, PSII densities appeared elongated along their major axis, and agreement with the ground truth dropped substantially. We benchmarked rejection sampling against MemBrain-pick^16^ and MPicker^13^, which we trained on this dataset. Across all membranes, rejection sampling achieved a median F1 score of 0.82, compared with 0.77 for MemBrain-pick and 0.74 for MPicker (Fig. 3d).

In summary, rejection sampling with continuous constraints matched deep-learning-based pickers on this dataset while requiring only a reference structure, rather than annotated training data, as input.

### C. Rejection sampling recovers ATP synthase organization

Having validated rejection sampling on both synthetic and experimental data, we next assessed its practical utility for complex membrane morphologies by applying it to ATP synthase in primary rat hippocampal neurons. Neurons are among the most energy-demanding cells, relying on a continuous supply of ATP to sustain ion gradients, synaptic transmission, and axonal transport^23^. This demand is met by mitochondria, whose inner membrane forms an intricate network of cristae that dramatically increases the surface area available for oxidative phosphorylation^24^. The ATP synthase, the terminal enzyme of this process, assembles into dimers that further associate into rows along highly curved cristae ridges, where they contribute to membrane curvature and cristae morphology^25^.

Rejection sampling evaluates only positions admitted by the constraints, so gaps in the surface model translate directly into regions where no particle can be found. The mitochondrial membrane architecture of neurons ranged from narrow lamellae to tubular cristae, a diversity that no single segmentation model captured (Fig. S2). We therefore used two segmentation approaches and found that their combination generally improved coverage. More specifically, MemBrain-seg^16^ recovered lamellar sheets more completely, whereas TomoSegNet^26^ performed better on tubular segments. We obtained the final segmentation mask by combining the results of both approaches and curating it in Mosaic^27^ to retain mitochondrial cristae membranes, from which we computed meshes for constrained matching.

Cristae occupy only a small fraction of the tomographic volume (Fig. 4a), so most cross-correlation scores computed over it fall outside the translational constraints and are discarded. We therefore developed a decomposition procedure via recursive bisection that restricts matching to a minimal set of subvolumes that cover the region of interest (Fig. S3). This avoided computing scores at inadmissible positions, reducing runtime by at least 2-fold for more than half of the dataset (Fig. S4a). Applying the constraints suppressed background cross-correlations (Figs. S4b and S4c), reducing the number of picks incompatible with the membrane geometry by 88% compared to unconstrained matching (Fig. S4d).

**Fig. 4.**
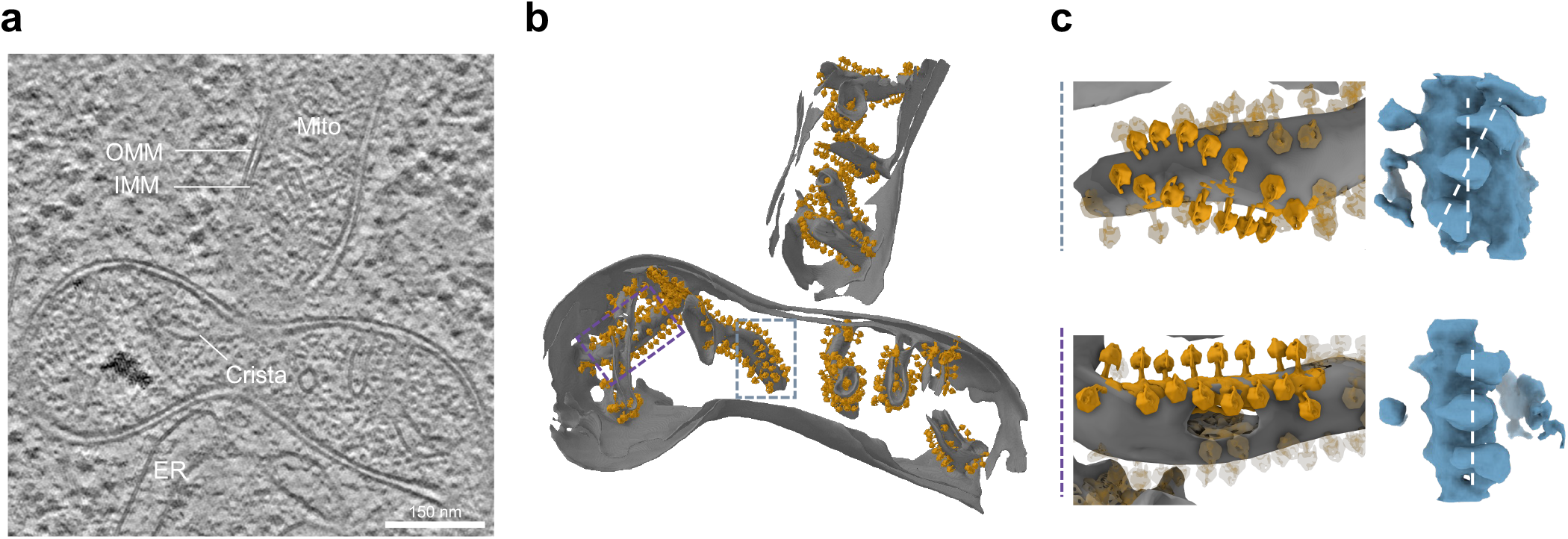
Rejection sampling localizes ATP synthase on mitochondrial cristae in primary rat hippocampal neurons. **a**, Tomographic slice through a primary rat neuron. Mito, mitochondrion; ER, endoplasmic reticulum; OMM, outer mitochondrial membrane; IMM, inner mitochondrial membrane. **b**, Mitochondrial membrane meshes (grey) with ATP synthase picks backmapped at the position and orientation assigned by rejection sampling (gold, shown is the template used for picking). Dashed boxes indicate the regions shown in (**c**), colored by inset. **c**, Helical (top) and linear (bottom) dimer rows on tubular cristae, with picks outside the respective region shown faded. Right, average densities of the picks in each row without further refinement, shown as isosurfaces. The vertical dashed line marks the tube axis as a common reference; the additional line in the upper row indicates the offset of the two heads. Enclosed dimer angles are 76.6° *±* 8.0° (helical, *n* = 14) and 79.1° *±* 6.0° (linear, *n* = 16; mean *±* standard deviation).

Mapping ATP synthase monomer picks back onto cristae surfaces, we found dimer rows on both lamellar cristae ridges and tubular cristae. These were predominantly linear, consistent with descriptions of mammalian mitochondria (Fig. 4b)^25^. In one instance, a row on a twisted tubular crista instead traced a helical course (Fig. 4c). Helical rows have also been described in *Polytomella*^28^, where they run along the ridges of disk-like cristae, but the combination with tubular cristae here is reminiscent of *Paramecium*^29^ and notable given the substantial difference in ATP synthase structure. While the co-occurrence of linear and helical rows is atypical within the same species, local averages were consistent with these assignments, and dimer angles fell within the range reported for mammalian and fungal ATP synthase (Fig. 4c)^30,31^. Whether helical rows occur reproducibly in mammalian mitochondria, and under which conditions, cannot be established from the 43 tomograms considered here. We therefore regard this as a hypothesis for future work on larger datasets and as a demonstration that our approach can surface the organizational detail such studies require in complex systems in situ.

## III. DISCUSSION

Identifying macromolecules in cryo-ET data remains challenging owing to low SNR, the missing wedge, and crowded backgrounds. Rejection sampling improves detection under these conditions by incorporating prior knowledge of macromolecular arrangement, thereby restricting template matching to biologically plausible translational and orientational configurations at voxel resolution with negligible computational cost. On synthetic IAV data, it improved detection, orientational assignment, and discrimination between macromolecules; on experimental spinach thylakoid data, it matched the performance of deep learning-based pickers informed by membrane structure without requiring training data. Applied to ATP synthase in neuronal mitochondria, it resolved two distinct row architectures across morphologically diverse cristae: predominantly linear rows alongside one helical row. This combination has, to our knowledge, not been reported within a single species. Rejection sampling is therefore a suitable tool for targeted discovery when prior geometric knowledge is available.

We see two principal limitations relevant to our approach. First, it inherits shortcomings of template matching by construction, in which suboptimal templates, masks, or filter parameters can adversely affect detection performance. Rejection sampling filters cross-correlation scores based on prior knowledge, but nevertheless requires sufficient signal in the underlying data for detection. The PSII dataset illustrated this, where agreement with the ground truth dropped on low-contrast membrane sheets despite the constraints. This limit is set by the cross-correlation metric itself, a linear filter that is optimal only under specific noise assumptions. Deep-learning-based detectors are not bound by those assumptions and should, in principle, be capable of exceeding this ceiling. Our benchmark should therefore not be read as a general ranking, but as a demonstration that rejection sampling is directly useful for macromolecular detection and could further help realize the potential of learned detectors by supplying the particle positions and orientations needed to train them. Second, rejection sampling is bounded by the quality of the constraints and, hence, the surface representation from which they derive. Incomplete representations exclude regions that may contain the protein of interest from the search, and obtaining adequate coverage can require combining complementary segmentation approaches, as we showed for ATP synthase on mitochondrial cristae. Even combining approaches did not always yield complete coverage, though such gaps could be closed by mesh completion and surface modeling tools. Therefore, rejection sampling moves part of the detection problem upstream, onto the construction of the surface model.

Because the constraint predicate is general, stronger constraints are possible within the same framework. Here, we constrained matches using the membrane normal, leaving the in-plane rotation free. On curved membranes, the principal curvature directions provide an additional reference that can fix the in-plane angle of proteins oriented relative to the curvature, such as the ATP synthase dimers along cristae ridges. More generally, constraints based on a protein’s symmetry group could be incorporated in the same way, tightening the search wherever such prior knowledge is available.

In summary, rejection sampling improves detection by enforcing geometric constraints at voxel resolution while retaining the scalability of an unconstrained search. It thereby extends constrained template matching to large, highly curved membrane systems that pervade cells and represents a step toward routinely resolving their molecular organization in situ.

## IV. METHODS

### A. Theory

#### 1. Template matching

Template matching localizes particles of interest by evaluating the cross-correlation (CC) between a tomogram *f* and a template *g* over all translational and rotational degrees of freedom. For a translation ***t***, the CC is defined in real space and, by the convolution theorem, equivalently evaluated in Fourier space,

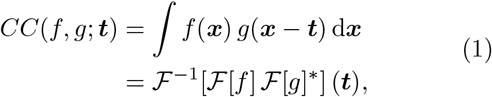

where ℱ and ℱ^−1^ denote the forward and inverse Fourier transform and ^*\**^ the complex conjugate. While a direct evaluation of the real-space integral scales as (*n*^2^) for *n* voxels, the Fourier-space formulation reduces the computational complexity to *O*(*n* log *n*).

For the template to be comparable to the tomogram, it must carry the same modulation imposed during imaging. We therefore apply the filter operator *W* to the template, defined as *W*[*g*] = ℱ ^−1^[*W* ℱ[*g*]], where the transfer function *W* combines the CTF with a tilt-dependent weighting that encodes the missing wedge.

Rotations are incorporated by sequentially evaluating the CC against the filtered, rotated template 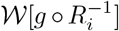 (The pullback 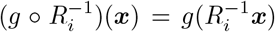 corresponds to an active rotation of the template, such that its reference axis *e*_*z*_ is mapped to *R*_*i*_*e*_*z*_), retaining the per-voxel maximum score together with the rotation producing it

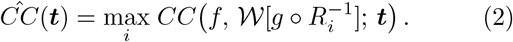

To account for changes in local brightness of the tomogram, the CC can be standardized by a background CC distribution derived from a template mask. Several formulations exist for this normalization; here, we use the fast local correlation (FLC, 32).

### **2**. Constrained matching using rejection sampling

Template matching is most efficiently performed in Fourier space, while translational and orientational constraints are naturally expressed in real space and lack an immediate Fourier-domain equivalent. Rejection sampling circumvents this by template matching in Fourier space and subsequently rejecting CC scores that do not satisfy the constraints in real space.

Constraints are encoded as seed points, e.g., sampled from a membrane surface, each defined by a position ***µ*** *∈* ℝ^3^ and a normal vector *n ∈* R^3^, which define a joint constraint predicate *C*(*R*_*i*_, *t*) that determines whether a match at translation *t* and rotation *R*_*i*_ is retained. Incorporating *C* generalizes Eq. (2) to

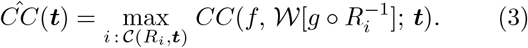

We define *C*(*R*_*i*_, ***t***) through a local coordinate system constructed from ***µ*** and ***n***. Specifically, we define a rotation matrix *R* that maps the default template orientation, chosen without loss of generality as ***e***_*z*_, onto ***n***, i.e., ***n*** = *R****e***_*z*_, and use it to generate an orthonormal basis around ***n*** ≡ ***n***_*z*_

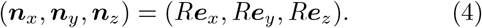

The orientational component of *C*(*R*_*i*_, ***t***) is satisfied if the effective template orientation ***r***_*i*_ = *R*_*i*_***e***_*z*_ falls within a cone of aperture *θ* around ***n***_*z*_, which in the local coordinate basis reads

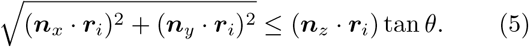

The translational component of *C* (*R*_*i*_, ***t***) is satisfied if ***t*** falls within an ellipsoid centered at ***µ*** and principal axes aligned along (***n***_*x*_, ***n***_*y*_, ***n***_*z*_)

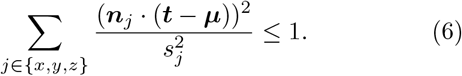

Here, *s*_*j*_ denotes the acceptance radius along ***n***_*j*_, but the spatial constraint generalizes to arbitrary masks.

### 3. From discrete to continuous constraints

In the discrete formulation, a given position can be covered by the masks of multiple seed points, introducing ambiguity in the orientational constraint. The constraint predicate *C*(*R*_*i*_, ***t***) extends naturally to a continuous formulation. By replacing seed points with a continuous normal vector field, we eliminate the effects of seed point spacing and orientational ambiguity. Further, this reduces the translation constraint to a single parameter *h*; the maximum distance to the surface. The continuous normal vector field is computed from a signed distance function (SDF) constructed using a triangular mesh representation of the surface *M*. The SDF *ϕ* : ℝ^3^ →ℝ is defined as

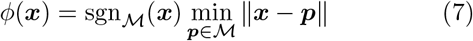

where sgn_*M*_ (***x***) ∈ {*−*1, +1} is an inside/outside indicator determined by the mesh winding number^33^. Since constraints are limited within *h*, SDF computation is restricted to |*ϕ*(***x***)| ≤ *h*, with query points determined via Euclidean distance transforms on voxelized meshes. The normal vector field ***n***(***x***) is then obtained from the gradient of the SDF

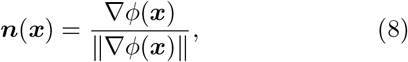

computed numerically on a discrete voxel grid using Gaussian-filtered finite differences (*σ* = 1, scipy v1.16.2, 34) to reduce noise and preserve gradient continuity.

Meshes derived from biological data are often imperfect. Therefore, we introduce three robustness measures. (1) The SDF sign definition becomes ambiguous in non-watertight meshes; this is addressed by optionally using precomputed vertex normals to determine sgn_*M*_(***x***) when mesh topology is unreliable. (2) The spatial band can extend laterally beyond open mesh edges, introducing spurious constraints. This is prevented by identifying boundary edges and rejecting voxels whose closest mesh point lies on such an edge and whose surface distance exceeds a suitably chosen cutoff (0 by default), determined via barycentric coordinates. (3) For closely apposed membrane sheets, such as thylakoid membrane stacks, the SDF gradient vanishes near the medial surface between adjacent membranes, producing unreliable normal vector estimates. Such voxels are instead assigned normals interpolated from the vertex normals of the closest mesh point via barycentric interpolation over the enclosing triangle. All mesh operations were implemented using Open3D (v0.19.0, 33).

Since constraints on individual voxels are independent in the continuous formulation, we can avoid an expensive scatter-add/scatter-or operation that was previously required to implement Eq. (6). This adaptation renders the computational cost of constraint enforcement negligible relative to that of the cross-correlation calculation.

### 4. Subvolume decomposition for sparse regions of interest

The ability of rejection sampling to operate on entire tomograms is a key computational advantage for broad regions of interest. On the other hand, it becomes increasingly inefficient as tomogram sparsity increases, as the majority of computed scores are rejected by the translation constraints. For such cases, we decompose the region of interest into a minimal set of subvolume covers that can be matched independently. We use the term subvolume to distinguish these regions from conventional subtomograms centered on individual particles. This decomposition is not specific to rejection sampling and applies equally to unconstrained template matching.

Let *U* ⊂ Z^3^ denote the region of interest, derived from seed point distributions or segmentation masks. We seek a collection of axis-aligned subvolumes ℬ = {*B*_1_, …, *B*_*N*_} covering *U* that minimizes the total computational cost

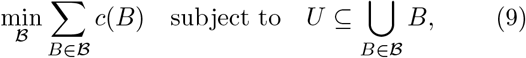

where 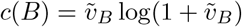 with 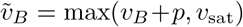 approximates the fast Fourier transform (FFT) cost of template matching against a subvolume of *v*_*B*_ voxels. The padding term *p* distinguishes the matching region from the effective FFT volume and reflects the boundary requirements of circular convolution. The saturation volume *v*_sat_ reflects the lower bound beyond which further reduction in subvolume size no longer reduces runtime, set by GPU utilization rather than the cost of the FFT itself.

We approach Eq. (9) through recursive bisection. Starting from the tight bounding box *B*_0_ ⊇ *U*, each box *B* with active voxels *X* = *B* ∩ *U* is partitioned along the axis-aligned plane that minimizes the combined cost of its children

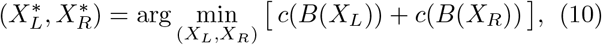

where (*X*_*L*_, *X*_*R*_) ranges over all bipartitions obtained by splitting between consecutive distinct coordinates along any axis, and *B*(·) denotes the bounding box of a voxel set. The recursion terminates at *B* when 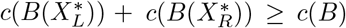, retaining *B* as a leaf; the resulting leaves form the cover. Sibling boxes can optionally be constrained to share a common shape during the recursion. The decomposition is accepted only if its total cost improves on that of the trivial bounding box over all voxels *B*_0_ by at least a factor of *τ* = 1.5 by default, and *B*_0_ is used otherwise.

Note that this procedure is a greedy heuristic restricted to nested binary partitions of the root bounding box; see Section XI B for a discussion of optimality.

### B. Data acquisition and processing

#### 1. Synthetic Influenza A data

Simulated IAV data were generated using Mepsi (v0.3, 19). The viral membrane was modeled as a sphere of radius 600 Å, with 387 HA and 85 NA instances (5:1 nominal target ratio before packing) placed at 10 nm spacing to reproduce experimental descriptions^17,18^. HA and NA templates were generated as previously described^27^. The system was solvated and converted into a tilt series ranging from 60° to −60° with 3° spacing, CTF-modulated at a defocus of 5 µm and SNR of 0.65, and reconstructed at a voxel size of 6.8 Å with an edge length of 300 voxels. Tomograms with different SNRs were obtained in the same way. Membranes were segmented using MemBrain-seg (v0.05, v10 alpha weights, 16) and processed in Mosaic (v1.1.4, 27).

#### 2. Spinach Photosystem II

Tomograms were downloaded from EMPIAR-12612^20^ at a voxel size of 14.08 Å. The corresponding ground truth annotations for PSII and membrane segmentations were obtained from doi:10.5281/zenodo.15090084^16,35^ and doi:10.5281/zenodo.15090120^20,36^.

#### 3. Rat neuron ATP synthase

Data were obtained from Schwarz *et al*. ^37^. Briefly, tomograms were processed in Warp (v2.0.0, 38), using AreTomo (v1.3.3, 39) for tilt-series alignment, and reconstructed at a voxel size of 10.73 Å. For membrane segmentation, we used MemBrain-seg (v10 beta weights) with an activation threshold of −1 on tomograms deconvolved using the provided utilities, and TomoSegNet (v1.2.0, model v4 weights, 26) on the Warp reconstructions. Segmentations from both tools were merged, curated, and clustered into cristae/non-cristae segments using Leiden clustering^40^ in Mosaic.

### C. Particle picking

#### 1. Template matching

All template matching was performed using PyTME (v0.3.4, 41). Templates were aligned to the z-axis, lowpass filtered to the Nyquist frequency of their respective tomograms at the voxel size used for template matching, and resampled to the corresponding voxel size using cubic-spline interpolation. For the IAV dataset, template structures were created as previously described^27^ at a box size of 60 voxels (408 Å). The PSII template was generated from EMD:6617^22^ with a box size of 48 voxels (675.84 Å). The ATP synthase template was derived using a combination of manual and automated picking in IMOD^42^ and GapStop^43,44^, respectively. False positives were removed by 3D classification in Relion, resulting in a 32 Å structure from 3,120 particles.

Template masks were adapted to the geometry of each template. For IAV, a cylindrical mask of outer radius 10 voxels (68 Å) and height 37 voxels (251.6 Å) with a sigma decay of 1 was centered on the template. For PSII, a cylindrical mask of radius 10 voxels (140.8 Å) and height 10 voxels (140.8 Å) with a sigma decay of 1 was centered on the template. The ATP synthase mask was generated by creating a density map from the PDB entries 5ARA, 5ARH, and 5FIK, which was downsampled to a voxel size of 10.73 Å. Using the mask creation utilities in Relion, the resulting map was extended by 2 voxels with a soft edge of 4 voxels^45^.

Seed points and surface meshes for rejection sampling were generated in Mosaic. For the IAV dataset, seed points were placed 80 Å radially outward from the mesh surface to approximate the center of mass of HA and NA instances. For the spinach dataset, membrane segmentations were converted to medial meshes via Poisson reconstruction of the medial skeleton, with normal vectors adapted to point in the expected orientation of PSII; this avoids conflicting normal-vector estimates that arise for meshes enclosing finite volumes when particles lie on the medial skeleton of the respective segmentation. For the rat neuron dataset, membranes were meshed using flying edges with windowed sinc smoothing (strength 80, 10 iterations, feature angle 120°), decimated tenfold by quadric edge collapse, followed by 10 iterations of Taubin smoothing.

Template matching was performed on AMD Epyc Genoa 9554 CPUs with NVIDIA L40S GPUs. Template matching and peak calling parameters for all datasets are summarized in Table 1. Corrections used generally follow from the data processing strategy. For spinach specifically, we did not use CTF or tilt weighting because it was unclear whether they would be reflected in the deposited denoised tomograms. Instead, we used a phase-scrambled template^43^ to correct for nonspecific background signal. `MaximumFilter` suppresses subsequent picks within a sphere of radius `min-distance` around each accepted peak, whereas `RecursiveMasking` suppresses the region covered by the template mask rotated into the orientation assigned during matching. The latter was used specifically for PSII, because an isotropic mask large enough to separate in-plane neighbors would also suppress genuine peaks on the apposed membrane sheet.

**TABLE 1.** Template matching parameters and peak calling strategies by dataset. Peak calling strategy parameters are specified as command-line flags in PyTME’s postprocessing module, without a double-dash prefix for brevity. Score cutoff based on n-false-pos. was computed as previously described^46^.

| Parameter/Data | IAV | Spinach | Rat |
| --- | --- | --- | --- |
| **Matching** |  |  |  |
| Score | FLC | FLC | FLC |
| Angular sampling [°] | 7 | 6 | 5 |
| Voxel size [Å] | 6.8 | 14.08 | 10.73 |
| Background correction | - | phase-scrambling | - |
| **Constraints** |  |  |  |
| Translation [voxel] | (6,6,10) and $h = 10$ | $h = 6$ | $h = 6$ |
| Orientation ($\theta$) [°] | 20 | 20 | 40 |
| **Corrections** |  |  |  |
| CTF | Multiply | - | Phase-Flip |
| Tilt weighting | - | - | Relion |
| **Peak calling** |  |  |  |
| Caller | Maximum-Filter | Recursive-Masking | Maximum-Filter |
| Strategy | mask-edges<br>num-peaks 650<br>min-distance 10 | mask-edges<br>num-peaks 3000 | mask-edges<br>num-peaks 4000<br>min-distance 6<br>n-false-pos. 5 |

#### 2. MemBrain-pick

MemBrain-pick (v0.0.8, 16) was trained on the spinach dataset. Training and validation sets were assembled by randomly selecting 3 tomograms each, with 5 membranes sampled per tomogram. Training tomograms were Tomo0022, Tomo0032, and Tomo0038; validation tomograms were Tomo0004, Tomo0010, and Tomo0017; and Tomo0001 was withheld as a held-out test set. The number of training membranes was determined by a data-efficiency analysis, which showed that detection performance remained flat beyond 9 training membranes^16^. Training was performed for 150 epochs and terminated upon convergence of validation loss at approximately 100 epochs (Fig. S5a). MemBrain-pick was trained on NVIDIA 4070 Super GPUs.

Peak calling parameters were determined by screening the mean shift bandwidth and the mean shift score threshold on the validation set. Bandwidth values from 60 to 120 and score thresholds from 1 to 12 were evaluated for average F1 score using step sizes of 5 and 1, respectively. A mean shift of 110 and a score threshold of 4 were identified as optimal.

#### 3. MPicker

Membrane meshes were generated from processed segmentations using MPicker’s meshing utilities (v1.2.0, 13) with a threshold of 4, and UV-unwrapped using Opt-Cuts^47^ with an energy of 4.04. Flattened tomograms were generated with a thickness of 10 and a radial basis function distance of 10. Picking was performed using EPicker (v1.1.2, 14), which we trained from scratch on the spinach dataset using the data split described above.

Training labels were derived by mapping the ground-truth particle coordinates into the flattened frame with MPicker’s coordinate conversion utilities, discarding particles further than 3 voxels from the reconstructed surface. Particles were labeled on the slice containing them and on one side of the slice. Flattened slices were padded with Gaussian noise to a square shape with edge length 1,024, following the procedure implemented in MPicker. This was done to bypass EPicker’s automated Fourier cropping, which we found corrupted training images smaller than that width. Padding also ensured consistent aspect ratios across slices of varying widths.

EPicker was trained for 150 epochs with default parameters: Adam with a batch size of 4 and a learning rate of 10^−4^, reduced tenfold at epochs 70 and 90. Since the MPicker interface does not allow custom validation datasets, we extended EPicker’s training loop to evaluate the loss on the validation set after each epoch, retaining the checkpoint with the lowest validation loss at epoch 36 (Fig. S5b). For each membrane in the held-out test set, the z-range containing the particles was identified by manual inspection. Picking was performed on the identified 3 to 4 voxel-thick slab, with a score threshold of 0, a maximum of 1,000 peaks, and a minimum peak distance of 7 voxels. Particle coordinates were converted into tomogram coordinates using the conversion utilities provided with MPicker. EPicker was trained and run on NVIDIA RTX 4070 Super GPUs.

#### 4. Comparison of approaches

Picks from all methods were evaluated against ground truth positions using a distance threshold of 6 voxels, allowing a maximum error of 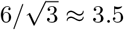 voxels per dimension, which corresponds to approximately half the radius of PSII and the IAV proteins at the respective voxel sizes. Each ground truth particle was uniquely assigned to the nearest pick. Angular deviation was quantified using the scalar product between the identified and ground-truth normal vectors.

A pick within the distance threshold of an assigned ground truth particle was counted as a true positive (TP), unmatched picks as false positives (FP), and unmatched ground truth particles as false negatives (FN). Precision and recall were computed as *TP/*(*TP* + *FP*) and *TP/*(*TP* + *FN*), respectively, and the F1 score as their harmonic mean. For the IAV dataset, detection performance was summarized by the area under the receiver operating characteristic curve, obtained by varying the score threshold over the called peaks.

For the spinach dataset, ground truth particles within 5 voxels of mesh boundaries were excluded from evaluation due to limited support in the tomographic densities (Fig. S5c), and the same mask was applied to pre-filter picks from all methods. The comparison is therefore restricted to regions where the ground truth is reliable, and does not resolve differences in performance at membrane boundaries. Template matching identifies particle positions based on center-of-mass correspondence with the reference template, whereas MemBrain-pick and MPicker detect salient structural features that do not necessarily coincide with the center of mass. These differences result in systematic offsets along the membrane normal, with rejection sampling picks localizing approximately 30 Å from the membrane surface envelope, while MemBrain-pick and MPicker picks localize at the surface envelope itself (Fig. S5d). To account for these baseline offsets, ground truth positions were projected onto the medial mesh for comparison with rejection sampling picks, while MemBrain-pick and MPicker picks were evaluated against the unprojected ground truth they were trained on. As shown in Fig. S5d, rejection sampling picks are better aligned with the projected ground truth in the medial plane, at a distance comparable to that of MemBrain-pick and MPicker picks relative to the unprojected ground truth.

Runtime was compared among unconstrained template matching, rejection sampling, and subtomogram-based matching by evaluating rejection sampling at seed point spacings ranging from 32 voxels to continuous constraints, with sequential halving intervals. Both approaches sampled SO(3) at 7° angular sampling density. Subtomogram-based runtime was approximated using 60-voxel subtomograms, sufficient to accommodate rotations while avoiding CTF aliasing, with cone-restricted rotational sampling at 7° angular sampling density, excluding redundant rotations arising from the C3 symmetry of HA. Time to evaluate a single rotation was subtracted from the total runtime, assuming cached imaging filters and FFT plans. All measurements were averaged across three independent runs.

### D. Data analysis

Three-dimensional renders of meshes, tomograms, and assembled molecular systems were created in Mosaic. Data analysis was performed in R (v4.5.0, 48) using ggplot2 (v3.5.2, 49) and data.table (v1.17.0, 50).

## V. ACKNOWLEDGEMENTS

We thank Andre Schwarz from the Max Planck Institute for Brain Research (Department of Synaptic Plasticity) for providing us with the primary rat hippocampal neuron dataset. We also thank the EMBL IT and HPC department for providing essential computational infrastructure^51^.

## VI. FUNDING STATEMENT

VJM and JK acknowledge funding from the CSSB flagship project, Plasmofraction, and from the European Research Council (ERC, TransFORM, 101119142). LD is a Minerva Fast Track Fellow in Erin Schuman’s department at the Max Planck Institute for Brain Research. The Schuman lab is supported by the Max Planck Society and the European Union (ERC Advanced Investigator Award Diverse Synapse 101054512 (EMS)). DMK acknowledges support by the German Research Foundation (DFG) via Germany’s Strategy-Cluster of Excellence Matter and Light for Quantum Computing (ML4Q, Project No. EXC 2004/1, Grant No. 390534769), individual Grant No. 508440990 and 531215165 (Research Unit “OPTIMAL”). LG acknowledges support from the Max Planck Institute for the Structure and Dynamics of Matter.

Views and opinions expressed are those of the authors only and do not necessarily reflect those of the European Union or the European Research Council Executive Agency. Neither the European Union nor the granting authority can be held responsible for them.

## VII. AUTHOR CONTRIBUTIONS

VJM conceived the method, implemented the software, performed the template matching experiments and analyses, and wrote the original draft. VJM and LG developed the theory. LD performed cryo-ET measurements and processing and contributed expertise on ATP synthase biology. DMK and JK acquired funding and supervised the project. All authors participated in reviewing and editing the manuscript.

## IX. CODE AVAILABILITY

All methods pertinent to rejection sampling described here have been implemented in PyTME and are available at https://github.com/KosinskiLab/pyTME. Constrained template matching is available starting with version 0.3.0 via the MaxScoreOverRotationsConstrained analyzer class, which is invoked at the level of individual rotations within PyTME’s user-defined analyzer architecture. The continuous SDF-based constraint formulation and the subvolume decomposition are available as dedicated modules from version 0.3.4. Analysis code and Mosaic sessions used throughout the study will be deposited in Zenodo.

## X. DATA AVAILABILITY

No data were collected in this study. Spinach data is available from Zenodo at doi:10.5281/zenodo.15090084^16,35^ and doi:10.5281/zenodo.15090120 ^20,36^. The primary rat hippocampal neuron data were acquired from Schwarz *et al*. ^37^.

## XI. SUPPLEMENTARY INFORMATION

### A. Angular resolution

Consider a parametric surface embedded in three-dimensional space described by *f*(***γ***) : *U* → *M* ⊂ ℝ^3^, ***γ*** = (*u, v*), where *u* and *v* are suitable coordinates ranging over the domain *U* ⊂ ℝ^2^. For a sphere, we can take spherical coordinates with (*u, v*) = (*θ, ϕ*) and *U* = [0, *π*] *×* [0, 2*π*] ⊂ ℝ^2^. A discretization of the surface *f* introduces an angular resolution for the surface normals ***n***(***γ***), which we illustrate in Fig. S6. It is directly related to the variation of the normal vectors along the surface. For a given direction d***γ*** = d*s* d***T***, with arclength d*s* and normalized tangent vector d***T*** ∈ *T*_***γ***_ *M*, we have the angular resolution

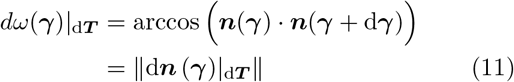

with d***n***(***γ***) _d***T***_ = ***n***(***γ*** + d***γ***) − ***n***(***γ***). A central result of differential geometry connects the normal vector variation to the shape operator *S*_***γ***_ : *T*_***γ***_*M* → *T*_***γ***_ *M* (also called Weingarten map) whose eigenvalues determine the local curvature of the surface^52^. Computationally it is represented as a two-dimensional matrix in the local tangent-basis (*∂*_*u*_*f, ∂*_*v*_*f*), where it depends on the first *I*_***γ***_ and second *II*_***γ***_ fundamental forms as 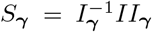. The angular resolution is hence given by

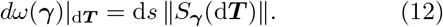

In the case of a spherical surface with radius *R*, this result simplifies to d*ω* = d*sR*^−1^, emphasizing that high curvature regions require dense sampling to retain angular resolution of surface normals.

### B. Optimality of subvolume decomposition

The greedy recursive bisection introduced in Section IV A 4 is a heuristic for the weighted set cover problem of Eq. (9). Each split is chosen on the immediate two-child cost, so the recursion cannot trade a locally suboptimal split for a globally better outcome, as the search is restricted to covers expressible as a binary sub-division tree of the root bounding box.

A strictly more general formulation casts the NP-hard set cover problem as an integer linear program (ILP) over a finite candidate set *S* of admissible subvolumes

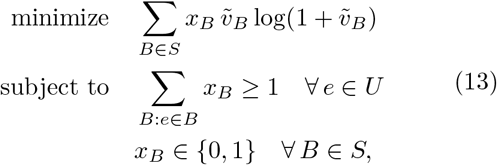

where each candidate *B* is defined by a position ***r***_*B*_ ∈ ℤ^3^ and an extent ***s***_*B*_ ℤ^3^, 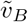 is the effective FFT volume defined in Section IV A 4, *x*_*B*_ is a binary selection variable, and *U* is the region of interest. Eq. (13) permits overlapping, non-nested, and arbitrarily structured covers, so an optimal ILP solution can yield a lower-cost cover than the greedy bisection.

In practice, this theoretical advantage is narrowed because the candidate set *S* must be enumerated in advance, restricting the search to a predetermined shape vocabulary (e.g., powers of two), whereas the greedy bisection generates tight bounding boxes adapted to the voxel distribution at each level. Empirically, we find that the greedy procedure produces covers of comparable total cost at substantially reduced solver runtime for the systems considered here.

---

**Fig. S1.**
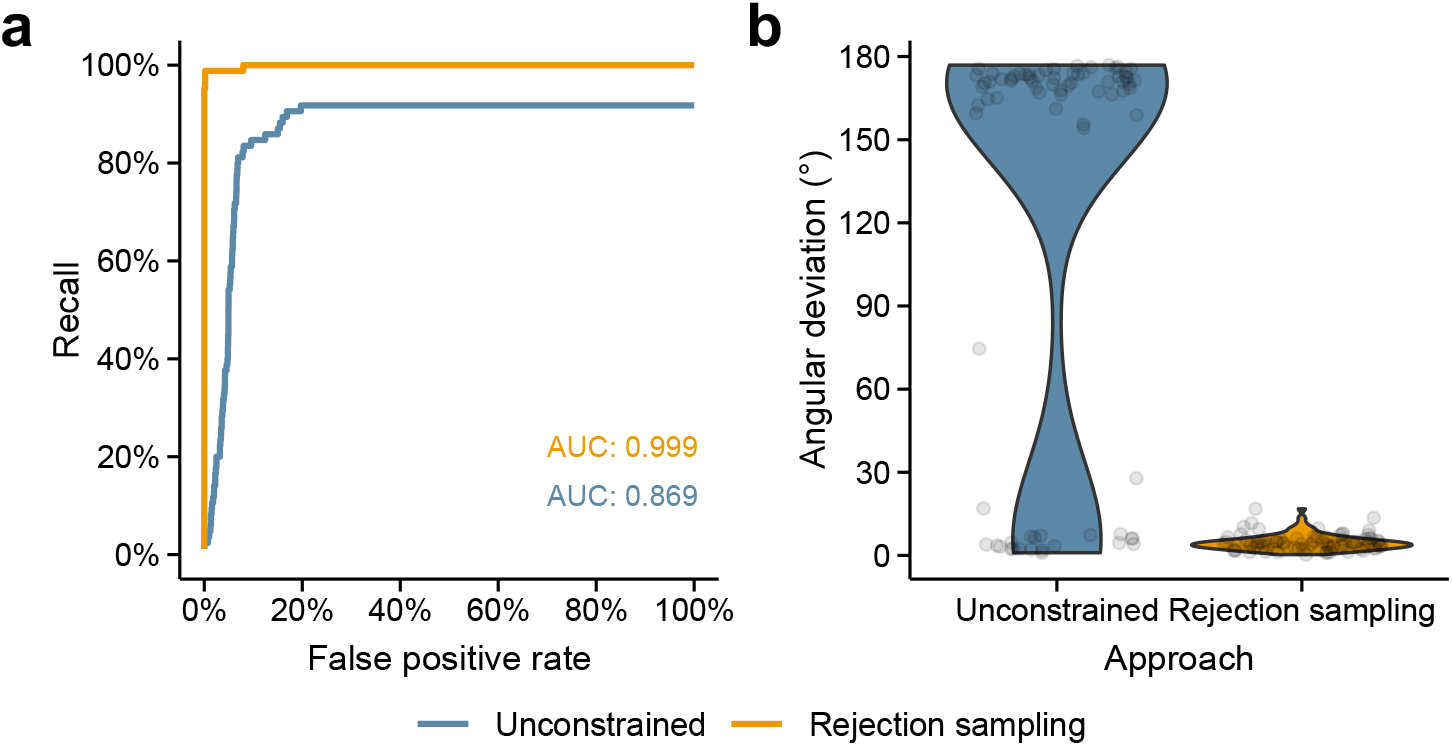
Comparison of template matching approaches on synthetic IAV data for the NA protein. **a**, ROC curve for NA detection and corresponding AUC values for each approach (top 650 picks). **b**, Angular deviation between matched and ground truth NA orientations (*n* = 85 and 78 for rejection sampling and unconstrained, respectively). Violins indicate kernel density estimates, and points indicate individual matches.

**Fig. S2.**
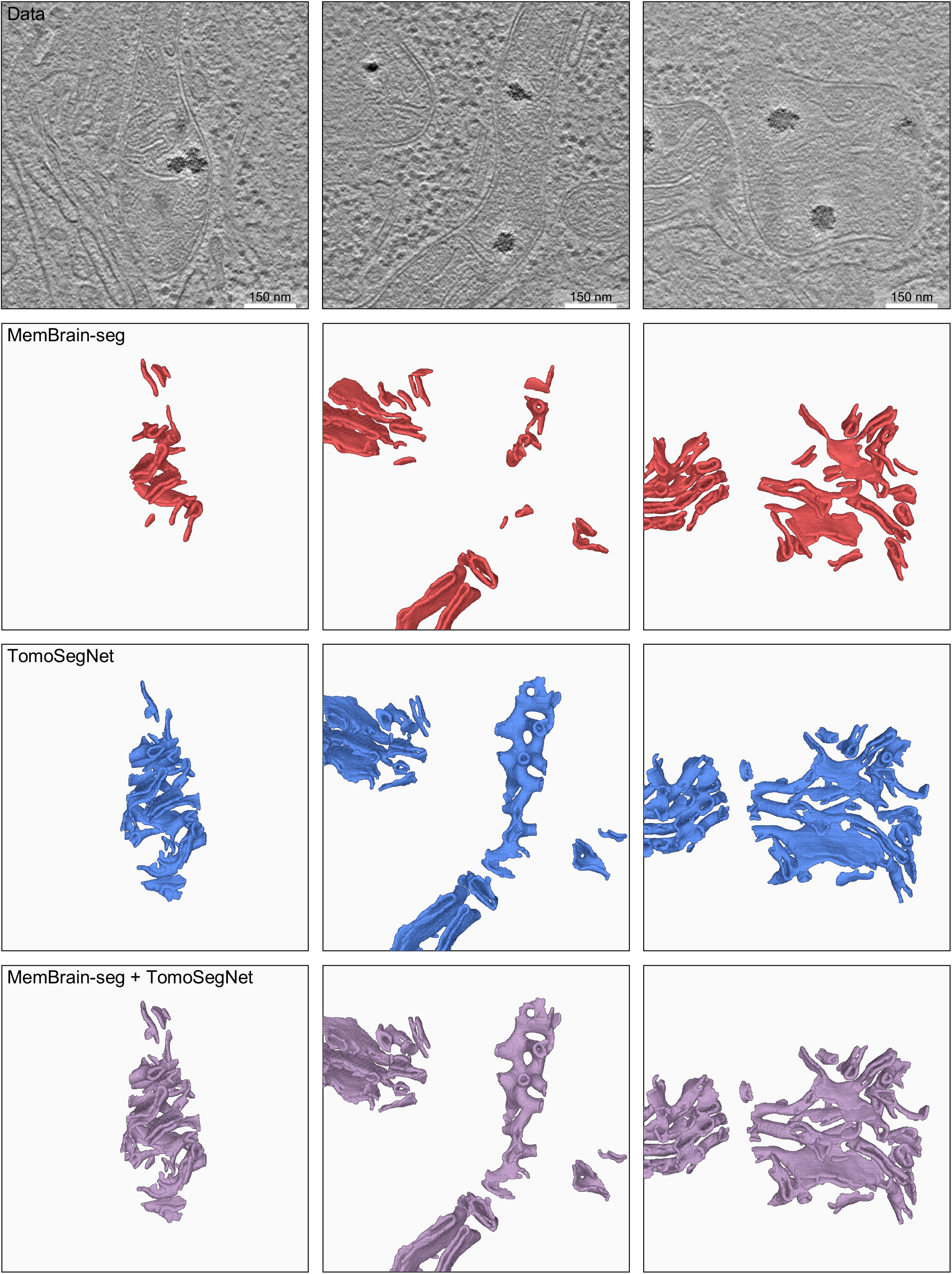
Rat neuron membrane segmentations obtained using MemBrain-seg and TomoSegNet. Segmentations were curated in Mosaic to exclude the outer mitochondrial membrane and non-cristae segments of the inner mitochondrial membrane.

**Fig. S3.**
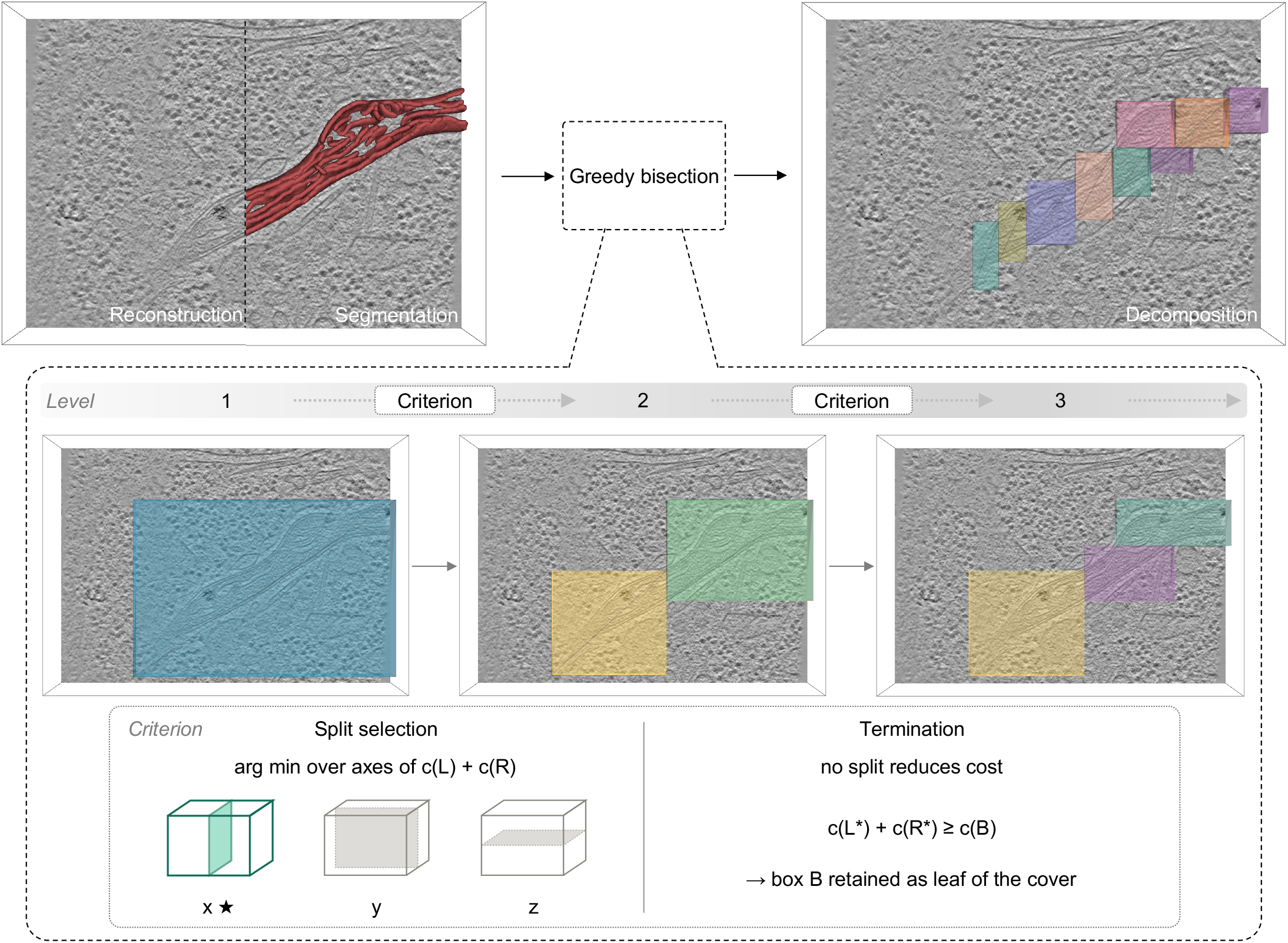
Illustration of subvolume decomposition using recursive bisection. A membrane segmentation (red) defines the region of interest *U* within the tomogram. Greedy bisection partitions *U* into a set of axis-aligned subvolumes (colored) that cover it and can be matched independently. The inset illustrates the recursion over three levels for a single membrane. Starting from the tight bounding box *B*_0_ ⊇ *U* (level 1), each box is split along the axis-aligned plane that minimizes the sum of the costs of its two children across all bipartitions (Eq. (10)). The recursion terminates when no split reduces the cost 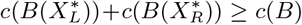.

**Fig. S4.**
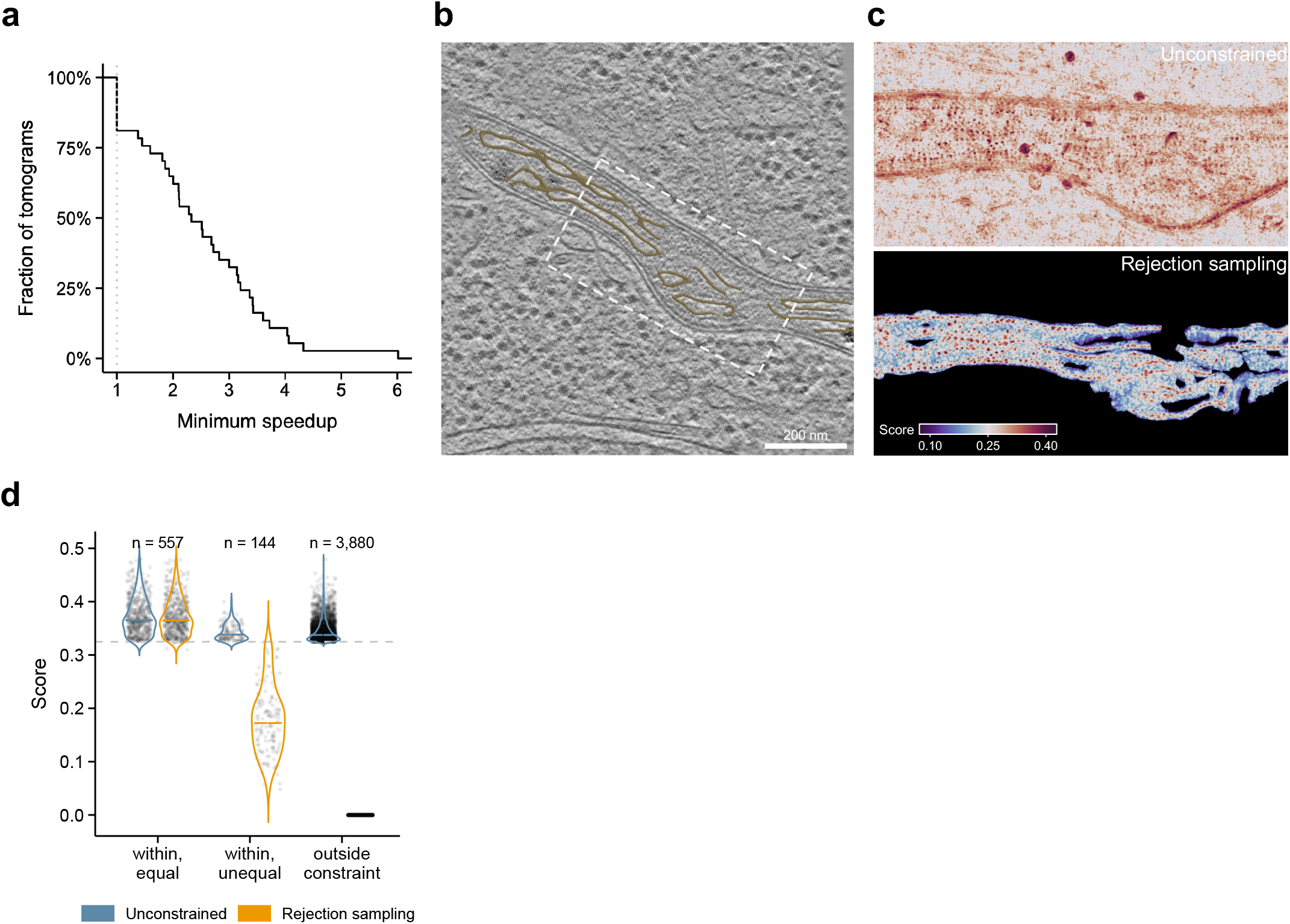
Supplementary analysis for ATP synthase localization. **a**, Runtime benefit of subvolume decomposition. Cumulative distribution of the speedup obtained by recursive bisection (decomposition) relative to matching a single bounding box around the segmentation mask (uniform, *B*_0_ in the notation of Section IV A 4). Speedup was computed as the ratio of wall-clock runtimes across the *n* = 43 tomograms of the rat neuron dataset. Tomograms at a speedup of 1 are those for which the minimum improvement was not met, and *B*_0_ was used instead. **b**, Tomographic slice of a mitochondrion in a primary rat hippocampal neuron with combined membrane segmentation overlaid (yellow). The dashed box marks the region shown in (**c**). **c**, Maximum intensity projection of cross-correlation score maps under unconstrained matching (top) and rejection sampling (bottom), on a shared color scale indicated in the bottom row. **d**, Template matching scores of picks above the significance threshold^46^ (dashed line, see Table 1) for unconstrained matching and rejection sampling, grouped by how the constraints affected them. Groups comprise picks within the membrane band whose score changed by 5% or less (within, equal), picks within the band whose score dropped by more than 5% (within, unequal), and picks outside the translational constraint (outside constraint). The score at a position is the maximum over the searched orientations, so restricting the admissible orientations can lower it. Positions outside the translational constraints are not considered by rejection sampling and are thus assigned a score of zero. Because the mitochondrial membrane segmentations appeared nearly complete, few genuine ATP synthase particles are expected outside the constraints, and the picks removed by the constraints therefore approximate the false-positive content of unconstrained matching. Violins indicate kernel density estimates, solid colored lines the median, and points individual picks.

**Fig. S5.**
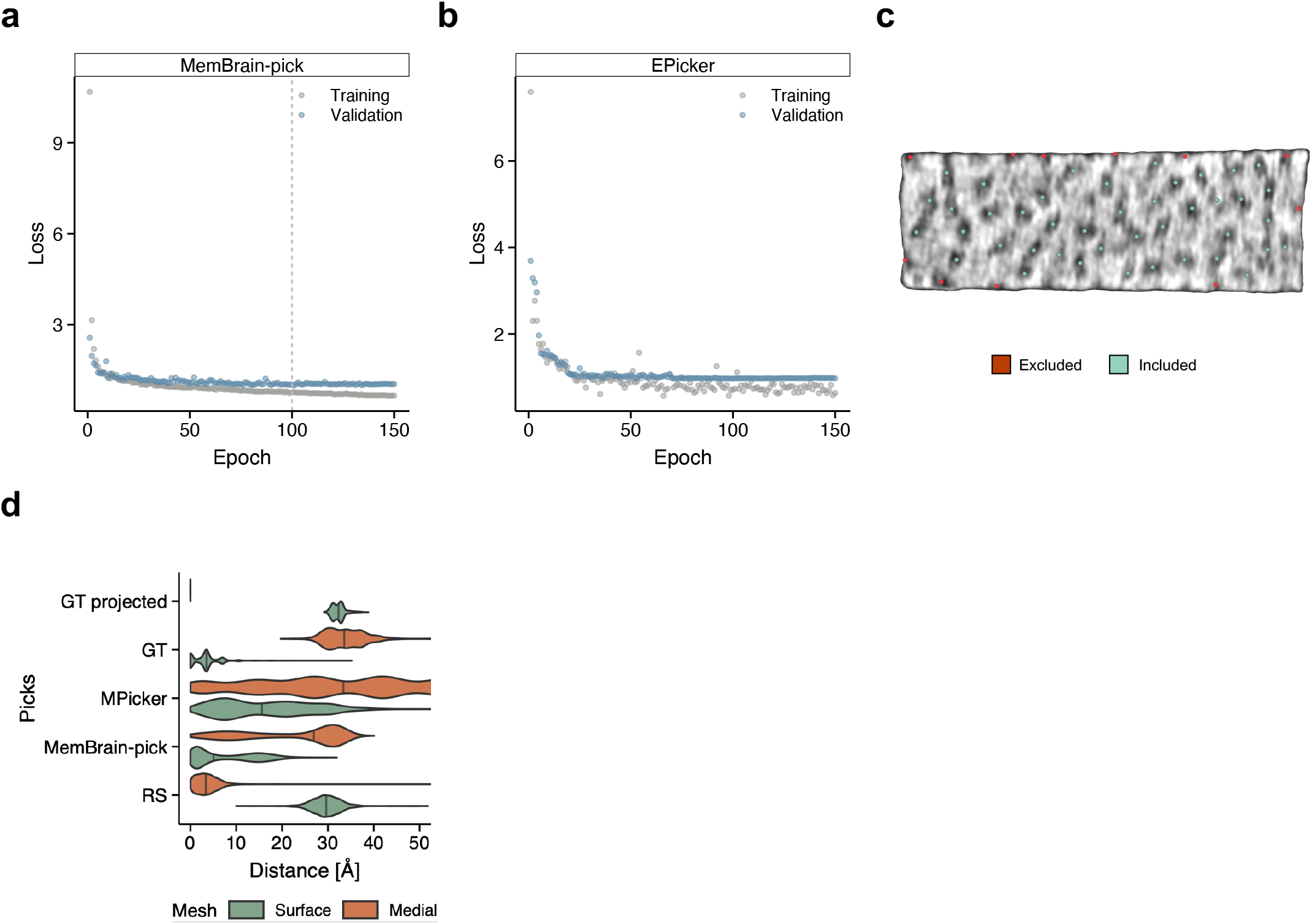
Supplementary analysis for PSII localization on spinach thylakoid membranes. **a**, Loss curves for MemBrain-pick training stratified by dataset. The dashed line marks the chosen model. **b**, Loss curves for EPicker training stratified by dataset. The dashed line marks the chosen model. **c**, Ground truth particles near membrane boundaries excluded from evaluation due to limited support in tomographic densities. **d**, Distance from the membrane surface for rejection sampling (RS), MemBrain-pick, MPicker, and ground truth (GT) picks, stratified by mesh type. RS picks localize to the medial plane consistent with the center of mass of PSII, whereas MemBrain-pick and MPicker picks localize to the membrane surface envelope, reflecting differences in the structural features each method is trained to detect. Violins represent the kernel density estimate, and solid lines represent the median.

**Fig. S6.**
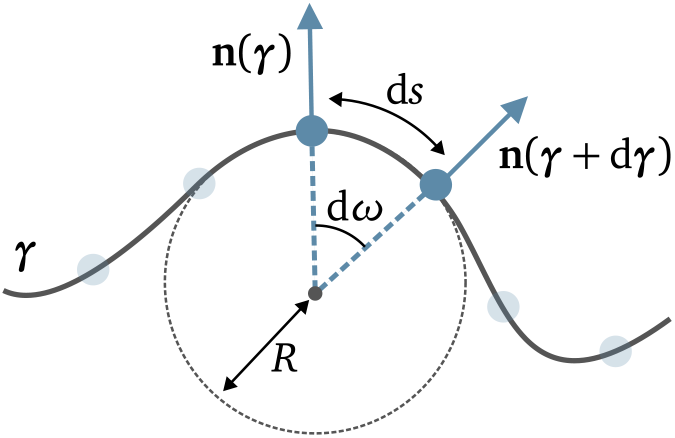
Illustration of surface normal angular resolution along a given direction d***γ*** = d*s*d***T***. The geodesic distance between sampling points d*s* (arc-length) and the local curvature of the surface, along the normalized tangent vector d***T***, here given by ∥*S*_***γ***_ (d***T***) = *R*^−1^, determine the angular resolution d*ω*|_d***γ***_ (Eq. (12)). This illustrates that high curvature regions require higher sampling rates to retain angular resolution.

